# Structural Acrobatics of IAA7: Exercising Fuzzy Regions to Aid Auxin-Mediated Catalysis

**DOI:** 10.1101/2025.06.08.657175

**Authors:** Jhonny Oscar Figueroa Parra, Maria Ott, Twinkle Bathia, Ryan Emenecker, Martin Wolff, Anja Thalhammer, Susanne Matschi, Luz Irina A. Calderón Villalobos

**Author notes:** To whom correspondence may be addressed: Luz Irina A. Calderón Villalobos and Jhonny Oscar Figueroa Parra; **Email:** and. **Author Contributions:** L.I.A.C.V., and J.O.F.P. designed research; J.O.F.P., M.O., T.B., R.E., M.W., A.T., and S.M. performed research; J.O.F.P., M.O., T.B., R.E., M.W., A.T., and S.M., analyzed data; and L.I.A.C.V., and J.O.F.P. wrote the paper. **Competing Interest Statement:** The authors declare no competing interests.

## Abstract

Auxin is the paragon of small molecules driving recognition of proteins for their ubiquitylation and degradation, enabling rapid decoding of external signals into transcriptional responses. Auxin functions as a molecular glue, enhancing interactions between the TRANSPORT INHIBITOR RESPONSE 1 (TIR1) and other TIR1-like proteins with various AUXIN INDUCIBLE/INDOLE-3-ACETIC ACID (AUX/IAA) transcriptional repressors. A Cullin RING E3 ubiquitin ligase, SCF^TIR1/AFBs^, mediates the ubiquitylation and degradation of AUX/IAA proteins by positioning them for ubiquitin transfer, leading to their rapid degradation by the 26S proteasome. AUX/IAAs contain a Phox Bem 1 (PB1) domain, a degron, and intrinsically disordered regions (IDRs) that influence auxin sensing and their polyubiquitylation by interacting with TIR1. However, the precise role of IDRs in AUX/IAA positioning for ubiquitylation, and the mechanism of multisite ubiquitylation within AUX/IAAs remain unclear. We used biophysical techniques and coarse-grained simulations (CGS) to study the structural behavior of IDRs in IAA7, a model AUX/IAA protein from *Arabidopsis thaliana*. Our findings show that IAA7 expands by approximately 2.6 nm upon binding to TIR1 in the presence of auxin, likely increasing the proximity between a lysine acceptor in IAA7 and the active site of the SCF^TIR1^ E3 ligase for ubiquitylation. Our data also suggest that IAA7 remains conformationally heterogeneous when bound to TIR1, forming numerous transient contacts along its IDRs and C-terminal PB1 with TIR1. We propose that these transient interactions outside the IAA7 degron interface enhance the TIR1·IAA7 association, resulting in highly dynamic and heterogeneous complexes that facilitate IAA7 multisite ubiquitylation, targeting it for rapid proteasomal degradation.

**Significance Statement:** In this study, we use biophysical techniques to elucidate the implications of AUX/IAA intrinsic disorder on their conformational ensemble and processing by the E3 ubiquitin ligase, SCF^TIR1^. Our findings demonstrate that IAA7 undergoes a structural expansion upon binding to TIR1 and remains highly dynamic, facilitating multisite ubiquitylation. These insights enhance our understanding of how ubiquitylation targets engage with their E3 receptor modules and underscore the structural advantages of intrinsic disorder in protein ubiquitylation targets.

## Introduction

A large percentage of eukaryotic proteins exhibit dynamic structural flexibility due to intrinsically disordered regions (IDRs) [1]. IDRs lack a well-defined three-dimensional (3D) structure under physiological conditions [2]. Instead, they can compact and extend, forming an ensemble of heterogeneous and dynamic conformations [3]. This flexibility allows IDRs to participate in protein-protein interactions through various mechanisms, including local patterning of aromatic residues, folding upon binding, or exposing short linear binding motifs (SLiMs) [4, 5]. These interactions significantly influence cellular signaling pathways. IDRs may harbor multiple distinct subregions of interaction that regulate the assembly of protein complexes, functioning as selective interaction hubs [6]. Given their flexibility, accessibility to enzymes, and diverse amino acid composition, IDRs are prime sites for post-translational modifications, including ubiquitylation [7].

Ubiquitylation enables the selective degradation of eukaryotic proteins through 26S proteasome-mediated proteolysis [8, 9]. This involves attaching ubiquitin molecules to a protein substrate, marking it for destruction. Ubiquitylation begins with the activation of ubiquitin by an E1 ubiquitin-activating enzyme (E1) in an ATP-dependent reaction, forming a high-energy thioester bond [10, 11]. Ubiquitin is then transferred from the E1 to an E2 ubiquitin-conjugating enzyme (E2) by a trans-thioesterification reaction [12, 13]. E3 ubiquitin ligases (E3), responsible for substrate specificity, bind to the E2 enzyme loaded with ubiquitin to facilitate the attachment of ubiquitin to the substrate [14, 15]. E3-ligase activity results in the formation of an isopeptide bond between the primary amino group on a lysine residue of the substrate and the carboxyl group of the ubiquitin moiety [16]. The specificity of this process is determined by the substrate-binding modules within the E3 ligases, which often recognize SLiMs that function as degradation signals (degrons) within the substrates [17, 18]. Degrons and specific ubiquitin chains flag the target protein for ubiquitylation and subsequent degradation by the proteasome [14]. Many degrons are flanked by IDRs, which are assumed to increase protein flexibility and enhance the likelihood of lysine residues accepting ubiquitin moieties [18]. However, the mechanism determining the location of ubiquitylation sites within IDRs remains unknown for many substrates.

In plants, auxin is a crucial small molecule for growth and development [19, 20]. Auxin enhances the recognition of AUX/IAA transcriptional repressors by a Cullin RING-type SKIP/CULLIN1/F-BOX PROTEIN (SCF)-E3 ligase, targeting them for degradation [21–23]. Nuclear auxin sensing occurs when an SCF-E3 ligase binds its ubiquitylation substrates, AUX/IAAs, in response to fluctuating auxin concentrations [21, 24]. Specifically, one molecule of auxin “glues” the substrate-binding domain of the F-BOX proteins TRANSPORT INHIBITOR RESPONSE 1/AUXIN F-BOX PROTEIN 1-5 (TIR1/AFB1-5) to a conserved degron motif in AUX/IAAs, allowing AUX/IAA polyubiquitylation and subsequent degradation by the 26S proteasome [21, 25, 26]. The degradation of AUX/IAAs is rapid, with half-lives between ∼6-80 min [25, 27, 28], enabling AUXIN RESPONSE FACTOR (ARF) transcription factors to regulate target genes that control various growth and developmental processes in plants, such as embryonic and post-embryonic development, tropisms, cell division, cell differentiation, and cell elongation [29–32].

The AUX/IAA degron acts as a substrate recognition site for SCF^TIR1^-mediated ubiquitylation in the presence of auxin [21]. The crystal structure of the auxin co-receptor system revealed that the 13-residue-long IAA7 degron motif adopts a single coiled conformation [21]. This conformation is stabilized by hydrophobic interactions with the auxin binding pocket in TIR1, positioning the core degron sequence GWPP of the IAA7 atop the auxin molecule [21]. Conservation of the AUX/IAA degron sequence is essential for SCF^TIR1^ recognition and subsequent degradation of AUX/IAA proteins. Mutations in the core degron sequence lead to AUX/IAA overaccumulation and constitutive repression of auxin-responsive genes by inhibiting ARF activity [30–33]. For instance, *axr2-1/iaa7* plants with a P87 to S substitution in the IAA7 core degron sequence accumulate IAA7 protein, resulting in repression of ARF activity [33, 34]. This causes gravitropic defects in roots, hypocotyls, and inflorescences, as well as reduced apical dominance in Arabidopsis plants [33, 34].

Additionally, AUX/IAA degrons are tripartite, comprising the E3 recognition motif (degron), lysine residues for ubiquitin transfer, and IDRs that are preferentially ubiquitylated [18, 26]. While the degron and the C-terminal Phox Beam 1 (PB1) domain are conserved among 23 of the 29 AUX/IAAs in *Arabidopsis thaliana* [35], the IDRs flanking the degron vary in length and amino acid composition, with different numbers of lysine residues as potential ubiquitin acceptors [36]. *In vitro* ubiquitylation assays have shown that IAA6, IAA7, IAA12, and IAA19 carry multiple ubiquitylation sites located in the degron-flanking IDRs, and the PB1 domain [26, 36]. Although the precise mechanism underlying this spread-out, multisite ubiquitylation pattern remains unknown, it may depend on the IDRs within AUX/IAAs and their binding mode to TIR1. However, due to the lack of a stable 3D structure, the functional implications of AUX/IAA intrinsic disorder for auxin sensing and ubiquitylation have remained elusive until recent years.

Crosslinking mass spectrometry (XL-MS) of ASK1·TIR1·auxin·IAA7 and ASK1·TIR1·auxin·IAA12 complexes revealed that the N-terminal IDR and C-terminal PB1 domains of IAA7 and IAA12 bind to opposite faces of TIR1 during auxin sensing [36]. Specifically, the PB1 domain binds to TIR1 at the leucine-rich repeat (LRR) regions 4 to 7, referred to as cluster 1 (residues 140 to 229), while the IDR upstream of the degron binds to cluster 2, consisting of LRR17–18 (residues 485 to 529) [36]. This suggests that the top surface of TIR1, separated by an approximate 7 nm distance between the two clusters, may accommodate an expanded conformation of IAA7 and IAA12 upon auxin-induced TIR1 binding. Nuclear magnetic resonance (NMR) studies and molecular dynamics simulations also demonstrated that residues outside the IAA17 degron motif make contact with the top surface of TIR1 [37]. Furthermore, coarse-grained simulations (CGS) and NMR data identified two predominant conformers of IAA17 when bound to TIR1 [37].

Despite recent models showing interactions between the IDR upstream of the degron and the PB1 domain of AUX/IAAs with TIR1, they provide a limited and static view of the conformational ensemble and flexibility of AUX/IAAs when bound to TIR1. A static conformation of AUX/IAAs bound to TIR1 does not account for the scattered ubiquitylation pattern of AUX/IAAs observed *in vitro* [26, 36]. Moreover, the active site of SCF-E3 ligases spans a confined 2.2 nm range, extending from the active site of SCF-E3 ligase to the lysine acceptor of ubiquitin [38]. Although some degree of flexibility has been observed for neddylated Cullin RING-type SCF-E3 ligases, the active form of Cullin RING ligases bound to NEDD8 [38] and interdomain linkers within F-BOX proteins for substrate positioning at the E3 active site [39], such a confined 2.2 nm active site range may rely on substrate flexibility. This flexibility may allow the target lysines within the substrate to approach the E3 active site, producing the multisite ubiquitylation observed for AUX/IAAs *in vitro* [38, 39].

In this study, we extend the understanding of the AUX/IAA and ASK1·TIR1 binding mode through various biophysical approaches coupled with CGS. Using IAA7 as a representative of the 29 AUX/IAAs in *Arabidopsis thaliana*, we demonstrate that its conformation expands upon binding to ASK1·TIR1 in the presence of auxin, likely increasing its proximity to the SCF^TIR1^ active site. Despite this expansion, IAA7 remains dynamic on top of TIR1. We show that IAA7 is conformationally heterogeneous while bound to ASK1·TIR1, forming multiple transient contacts with the top surface of TIR1. We propose that these multiple contact points enhance the affinity between ASK1·TIR1 and IAA7, thereby contributing to the stability of the complex. Additionally, our data suggest that the combination of IDR expansion and the conformational heterogeneity of IAA7 when bound to TIR1, facilitates its multisite ubiquitylation by SCF^TIR1^, targeting IAA7 for rapid proteasomal degradation.

## Results

### IAA7 exhibits low compactness and high conformational flexibility

The flexibility of E3 ligases is crucial for the proper orientation of target substrates at their active sites for ubiquitin transfer [38, 39]. However, ubiquitylation substrates such as AUX/IAAs possess IDRs, whose flexibility may also contribute to substrate orientation. The IDRs among the 23-degron-containing AUX/IAAs vary in length and may contribute differently to this process [36]. AUX/IAAs can be categorized by length into short (≤199 amino acids), medium (200-299 amino acids), and long (≥300 amino acids). For our biophysical studies, we selected IAA7 due to its medium length, which serves as a midpoint reference among the variously sized AUX/IAAs (**SI Appendix, Fig. S1**). To identify regions in IAA7 responsible for its conformational flexibility, we used dynamic light scattering (DLS) and static light scattering (SLS) to assess IAA7 compactness. DLS determined the hydrodynamic dimensions, represented by the Stokes radius (*R_S_*), while SLS ascertained the average oligomeric state (**SI Appendix, Fig. S2**). AUX/IAAs oligomerize via PB1-PB1 interaction due to a network of charged residues in a basic and an acidic patch on opposite faces of the PB1 domain [40–43]. To prevent oligomerization and aggregation, which could decrease protein solubility in our assays, lysine residues in IAA7 at positions 128, 138, and 139 were replaced with alanines. These positions correspond to the basic patch in the IAA7 PB1 domain, and the replacements were made for all our experiments [43]. Additionally, truncated variants of IAA7, containing only the N-terminal IDR or the C-terminal PB1, were expressed separately to assess their contribution to the overall compactness of IAA7. These variants were named IAA7^ΔPB1^ and IAA7^PB1^, respectively.

IDRs and globular proteins exhibit characteristic relationships between their *R_S_* and mass, described by a scaling law of type *R_S_=a×M^b^*, where *M* is the protein mass, and *a* and *b* are constants depending on the structural type of the protein [44]. This characteristic scaling behavior is visualized in an *R_S_* vs. mass plot (**Fig. 1A**). Globular proteins show a scaling exponent of *b*=0.33, while denatured proteins have a scaling exponent of *b*=0.493, resembling the disordered state. Both scaling exponents were determined from large protein sets [45]. Proteins with different degrees of folded domains and IDRs will show intermediate scaling behaviors between the globular and disordered limits, as they have a mixture of both structural types. The data indicates that IAA7 exhibits intermediate scaling behavior, suggesting it is neither completely folded nor entirely disordered (**Fig. 1A**). To identify the regions in IAA7 corresponding to each structural type, we analyzed the split IAA7 variants, IAA7^ΔPB1^ and IAA7^PB1^. This allowed us to determine the contribution of each protein half to the overall structure of IAA7. Our findings revealed that IAA7^ΔPB1^ scaled closer to disordered structures, while IAA7^PB1^ resembles globular proteins more closely than the full-length IAA7 and IAA7^ΔPB1^ (**Fig. 1A**). This suggests that IAA7 consists of an IDR and a folded domain, which aligns with disorder predictions and previously published structural data [36].

**Fig. 1.**
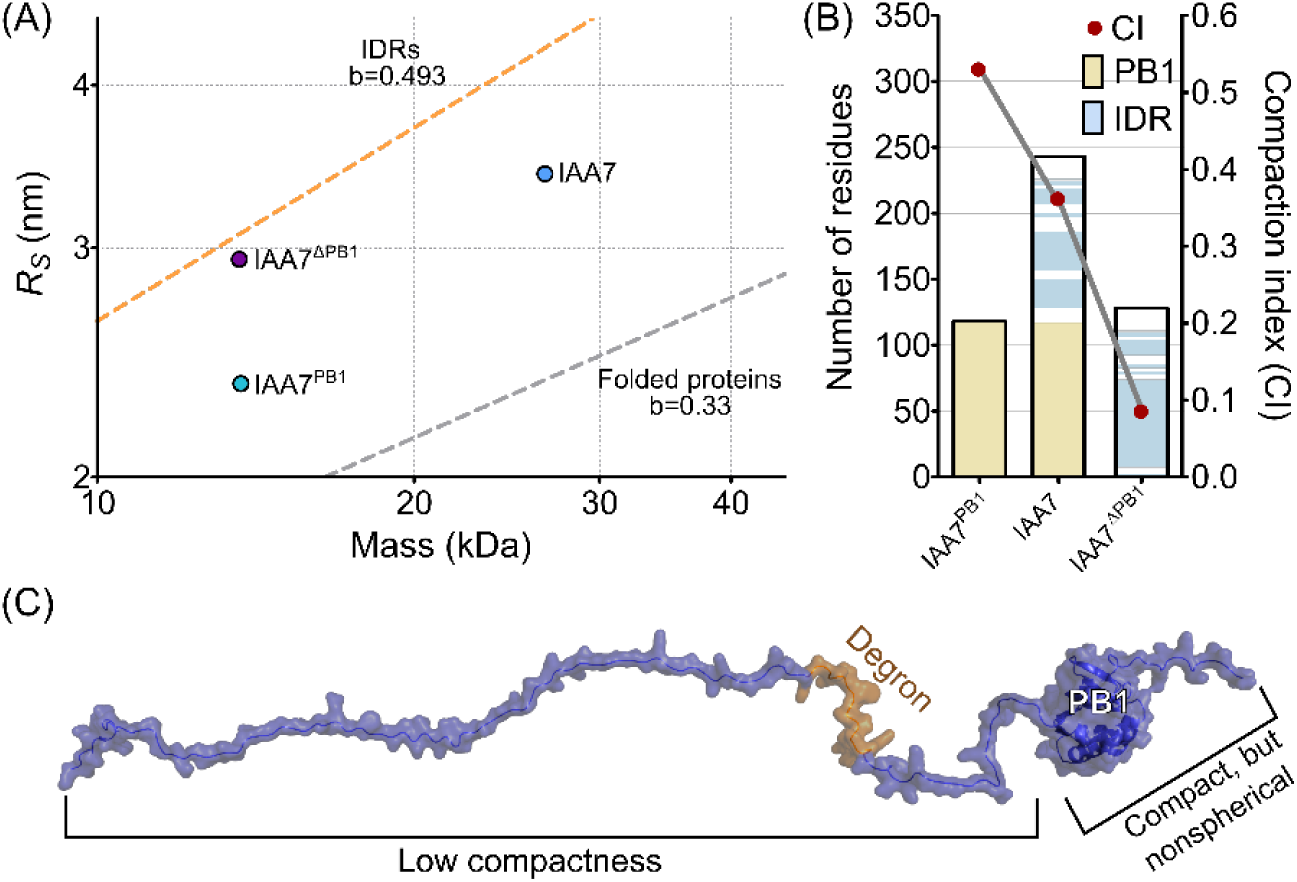
IAA7 exhibits low compactness. (A) Stokes radius (*R_S_*) vs. mass plot of IAA7 variants. The dashed lines represent the scaling behavior for IDRs (orange) and globular proteins (gray). The b values show the scaling exponent in the equation *R_S_ = a*M^b^*, taken from [45, 60]. (B) IAA7 sequence in bar representation showing the PB1 (yellow) and IDR with IUPRED disorder scores ≥0.5 (https://iupred2a.elte.hu/) (blue). The compactness, expressed as CI, of IAA7 decreases linearly in the following order: IAA7^PB1^>IAA7>IAA7^ΔPB1^. (C) AlphaFold2 model of IAA7 shows the degron location in a low compactness region at the N-terminus, while the PB1 domain at the C-terminus is more compact but has a nonspherical shape due to a disordered tail.

To interpret the scaling behavior of IAA7, we used *R_S_* values to determine the compaction index (CI). The CI provides a relative measure of protein compactness by comparing the scaling exponents of disordered and globular proteins relative to their mass [46, 47]. In globular proteins, CI equals 1, whereas in disordered proteins, CI equals 0 [46, 47]. Among the three variants, IAA7^PB1^ showed the highest compactness. However, IAA7^PB1^ (CI = 0.53) exhibits lower compactness compared to typical globular and spherical proteins, likely due to a short C-terminal IDR at the end of the PB1, as identified by various structural predictions [36] (**Fig. 1B, C**). A low compactness characteristic of disordered states was observed in IAA7^ΔPB1^ (CI = 0.08), suggesting greater conformational flexibility in the N-terminal side of IAA7 compared to its PB1 domain (**Fig. 1B, C**).

We further complemented these findings with small-angle X-ray scattering (SAXS) coupled with an ensemble-optimized method (EOM) to analyze IAA7 (**SI Appendix, Fig. S3**). EOM, which interprets SAXS patterns by considering a range of conformations, provides a more accurate representation of the IAA7 pervaded volume [48]. Using EOM, we generated a pool of 10,000 random conformations, computed theoretical SAXS profiles, and determined their radius of gyration (*R_g_*). The *R_g_* measures the distance between the protein’s center of mass and its outer regions, serving as a proxy for protein compactness [49]. The best-fitting SAXS profiles were described by a subset of the ensemble with a mean 〈*R_g_*〉=3.5 ± 0.2 nm (n = 2). This selected pool suggests that IAA7 adopts conformations that are more expanded than a globular protein but not as extended as a fully disordered chain lacking internal interactions (**Fig. 2A**). The selected IAA7 pool also shows an extended tail due to the variety of IAA7 expanded conformations in solution (**Fig. 2A**) [48]. In addition, we compared our experimental SAXS data to simulated SAXS values generated from conformations obtained from CGS using the mPiPi model one-bead-per-residue representation of IAA7 [50]. We used autoSCTR [51] to generate SAXS values based on our simulations and then rescaled the simulated SAXS values to compare them to the experimental values. We found that the simulated SAXS values were very similar to our experimental SAXS of IAA7 (**SI Appendix, Fig. S4**). From our CGS we determined the 〈*R_g_*〉 = 3.8±0.1 nm (n=3), which is consistent with our EOM analysis (**Fig. 2B**). Next, we took conformations from frames that had *R_g_* values evenly spaced between our most compact and most expanded conformations to visualize the range of ensembles that IAA7 adopts (**Fig. 2C**). The broad conformational diversity of IAA7, spanning from compact to expanded states, is attributed to its N-terminal IDR. Taken together, the results show that IAA7 has low compactness and explores multiple conformations due to its IDR.

**Fig. 2.**
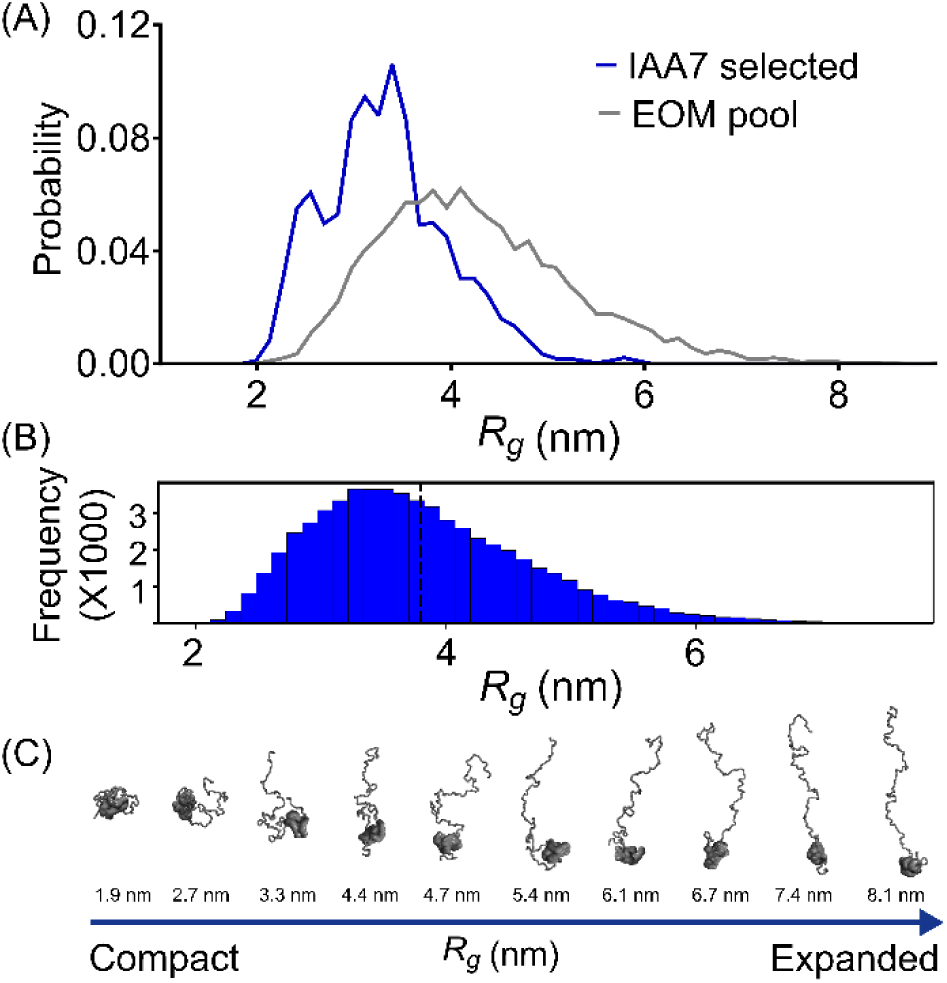
IAA7 displays IDR-characteristic expanded conformations. (A) EOM for IAA7 shows the *R_g_* distribution of the ensemble pool (gray) and the selected IAA7 ensembles (blue). (B) Frequency distribution of IAA7 *R_g_*from CGS, with the black dashed line representing the mean of the distribution. (C) Snapshots of IAA7 conformations identified by CGS reveal a broad spectrum, ranging from compact to expanded structures.

### IAA7 expands upon binding to TIR1

Based on SAXS and CGS results, we determined that IAA7 adopts multiple conformations in its unbound state, which are more expanded than those of a typical globular protein. Niemeyer et al. (35), suggest that an expanded conformation of IAA7 facilitates its binding to TIR1 at three distinct interfaces, forming the ASK1·TIR1·auxin·IAA7 receptor system. Specifically, the degron at the center of IAA7 locks auxin into the TIR1 auxin-binding pocket, the PB1 domain binds to TIR1 cluster I, and the IDR, located downstream of the degron, interacts with TIR1 cluster II. Given that TIR1 clusters I and II are approximately 7 nm apart in the 3D structure, we hypothesized that the IAA7 conformational ensemble average expands when fully engaged with TIR1 binding (**Fig. 3A**).

**Fig. 3.**
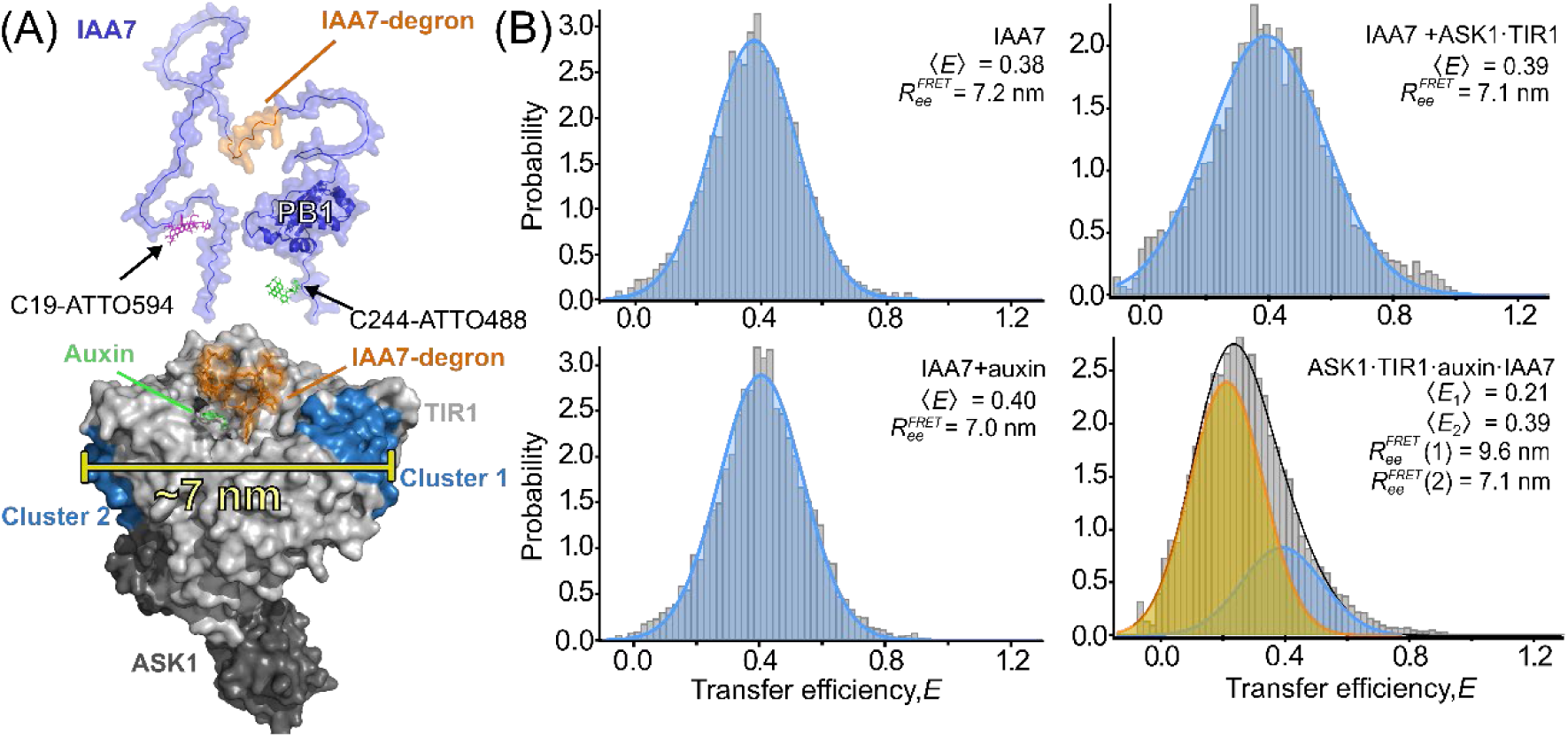
smFRET analysis shows IAA7 expands upon auxin-induced ASK1·TIR1 interactions. (A) The AlphaFold2 model of IAA7 highlights modifications at C19 with ATTO594 (magenta) and at C244 with ATTO488 (green). Modification of C19 with ATTO488 and C244 with ATTO594 was also possible due to labeling stochasticity but is not depicted here for simplicity. The TIR1 (PDB: 2P1Q) cluster 1 and 2 spaced by a 7 nm distance bind IAA7 PB1 and IDR independently of auxin, respectively. (B) Transfer efficiency histograms of IAA7 (50-100 pM) labeled with ATTO488/594 in the presence of auxin (indole-3-acetic acid, 10 µM) or ASK1·TIR1 (1 µM), or both. Blue lines represent the Gaussian fit of unbound IAA7, the orange line represents IAA7 bound to ASK1·TIR1, and the black line is the sum of the distributions. smFRET measurements were performed in three independent replicates, and transfer efficiency bursts were combined into a single file for analysis.

To test whether IAA7 expands when binding to ASK1·TIR1, we performed single-molecule Förster resonance energy transfer (smFRET) experiments. smFRET measures the energy transfer efficiency (*E*) between two adjacent fluorescent dyes, which is inversely proportional to the inter-dye distance. Using ATTO488/ATTO594 donor/acceptor pair dyes, IAA7 was labeled at the PB1 domain and the IDR, targeting the solvent-accessible cysteines at positions 19 and 244 (**Fig. 3A**). Fluorescence labeling was validated by in-gel fluorescence detection combined with LC-MS/MS to determine site-specific dye-IAA7 conjugates (**SI Appendix, Fig. S5-S6, Table S1**).

First, smFRET was performed on IAA7 without auxin or ASK1·TIR1. **Fig. 3B** shows the Gaussian fits of the *E* histograms of IAA7, where 〈*E*〉 denotes the mean value of the distribution. The 〈*E*〉 values were converted to ensemble and time-averaged distances, yielding inter-dye distances, referred to as 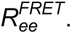 IAA7 showed an 〈*E*〉=0.38, equivalent to 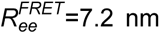 (**Fig. 3B**). Next, smFRET was performed with IAA7 preincubated with either auxin or ASK1·TIR1. Preincubation did not impact 〈*E*〉, resulting in 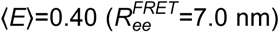 and 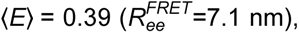 respectively (**Fig. 3B**). This indicates that neither auxin nor ASK1·TIR1 alone triggers a large change in the IAA7 conformational ensemble. However, we observed an increased width of the *E* distribution when IAA7 was incubated with ASK1·TIR1 (**SI Appendix, Table S2**), likely due to weak ASK1·TIR1 auxin-independent interaction as reported elsewhere [22, 36].

When we incubated ASK1·TIR1, IAA7, and auxin, to form an auxin receptor complex, 〈*E*〉 decreased to 0.21, corresponding to an 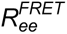 of 9.6 nm in 73% of the IAA7 molecules, indicating an expanded IAA7 conformation (**Fig. 3B**). The remaining 27% corresponded to an unbound or free IAA7 fraction with 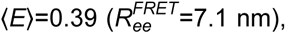 possibly due to spontaneous inactivation of the IAA7 degron, rendering it unable to bind ASK1·TIR1 effectively [37]. We concluded that TIR1 and auxin stabilize an expanded IAA7 conformation to form the auxin receptor complex.

### IAA7 remains dynamic when bound to ASK1·TIR1

After determining that the IAA7 conformational ensemble average expands when bound to TIR1, we next investigated whether this IAA7 conformation is static or dynamic. To assess the conformational dynamics of IAA7, we evaluated the distribution of inter-dye distances using fluorescence information by determining the relative donor lifetime in the presence of the acceptor, *τ_DA_*⁄*τ_D_*. This parameter provides information about the variance of the underlying distribution of transfer efficiencies [52]. Typically, for a protein with rigid or static conformation, the values of *τ_DA_*⁄*τ_D_* cluster close to the diagonal, corresponding to a single and fixed inter-dye distance [52]. However, IDRs exhibit a broad and rapidly sampled inter-dye distribution due to changes in their conformation, with values of *τ_DA_*⁄*τ_D_* clustering above the diagonal [52].

The lifetime analysis showed that the *τ_DA_*⁄*τ_D_* distribution of IAA7 alone or IAA7 with auxin clustered above the diagonal. This indicates that IAA7 is dynamic, and that auxin alone does not influence its conformational dynamics (**Fig. 4**). Similarly, the fluorescence lifetime analysis of IAA7 in the presence of ASK1·TIR1, or auxin and ASK1·TIR1 showed that IAA7 conformation remains dynamic, clustering above the static line (**Fig. 4**). Overall, this indicates that instead of adopting a single expanded conformation, IAA7 exhibits structural acrobatics on top of TIR1 and adopts an ensemble of multiple expanded conformations during auxin perception.

**Fig. 4.**
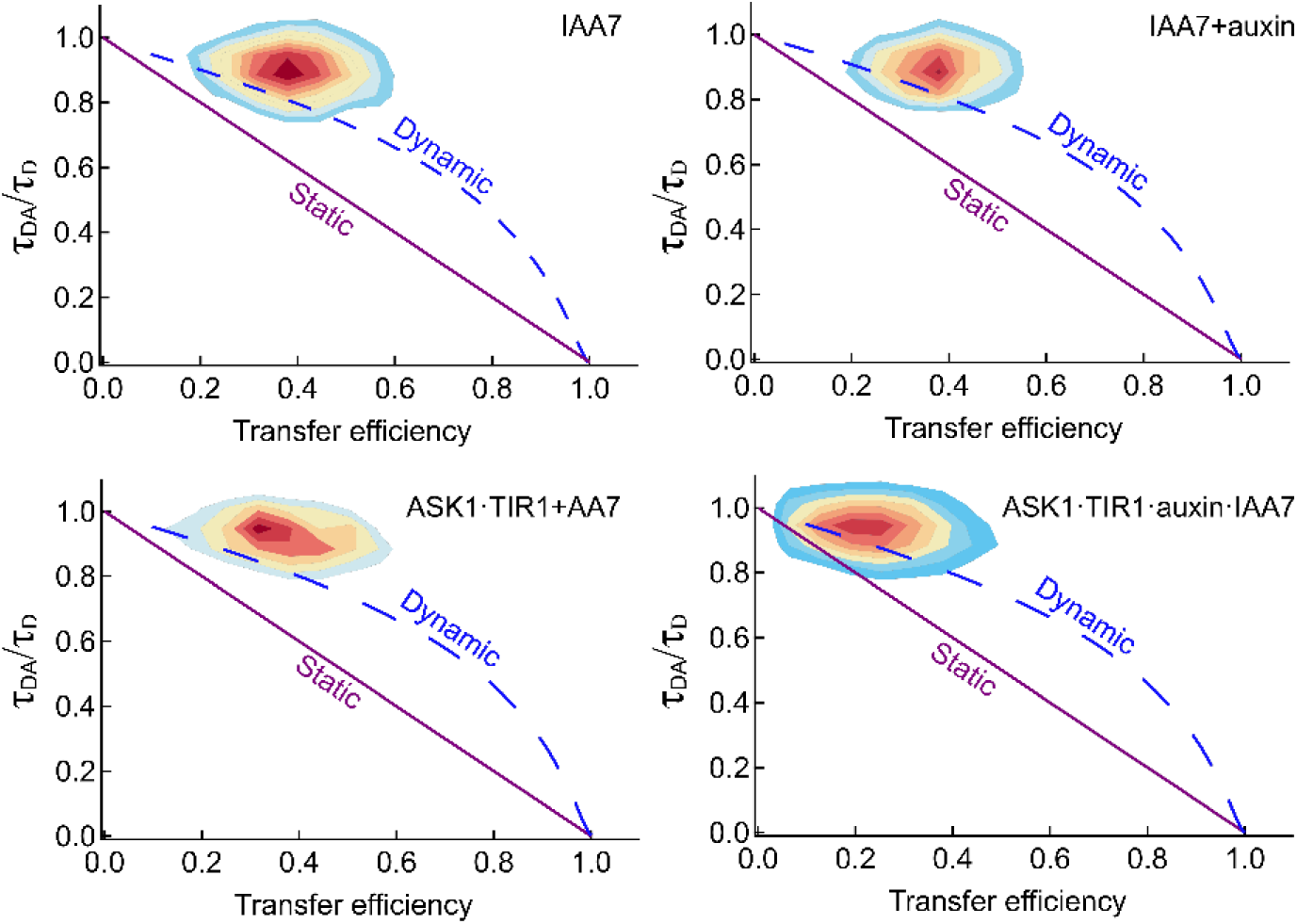
Fluorescence lifetime analysis shows the IAA7 conformational acrobatics. The 2D-correlation plot shows the relative donor fluorescence lifetimes in the presence of the acceptor (*τ_DA_*/*τ_D_*) with respect to the transfer efficiencies derived from the fluorescence intensities of the donor and the acceptor molecules. The static line represents a fixed donor-acceptor distance, the dynamic line results from conformational sampling using a self-avoiding walk (SAW) polymer model.

### IAA7 is conformationally heterogeneous during auxin perception

To complement the smFRET results and visualize the conformational dynamics of IAA7 when bound to TIR1, we performed CGS [50]. We hypothesized that CGS could enhance our molecular modeling and bridge our experimental techniques by providing a visualization of the dynamic auxin receptor complex [53]. First, we carried out simulations where IAA7 was not bound to ASK1·TIR1, representing an auxin-free state. For these auxin-free simulations, we aimed to identify residues within the IDR and PB1 domain of IAA7 that may drive interaction with ASK1·TIR1 in the absence of auxin. To reduce the computational cost, CGS was performed in a 500Å-by-500Å space (**SI Appendix, Fig.S7**). This confined space allowed us to determine IAA7 residues that contribute to ASK1·TIR1 interaction.

Interaction frequencies between IAA7 and ASK1·TIR1 were calculated based on the inter-protein residue contact frequency. For example, if an IAA7 residue was within 10 Å of an ASK1·TIR1 residue in every simulation frame, the contact frequency would be 100%. Conversely, if no contact occurred throughout the entire simulation, the contact frequency would be 0%. We found that the IAA7 core degron motif, GWPPV, and its immediate neighboring residues showed a high frequency of interaction with ASK1·TIR1, despite the absence of auxin in our simulations (**Fig. 5A**). These results align with other experimental evidence supporting a strong role for the degron motif in driving IAA7 interaction with ASK1·TIR1 independently of auxin [21]. Interestingly, residues near the GWPPV motif and at the PB1 domain in IAA7 exhibited higher interaction frequencies with ASK1·TIR1 than those driven by the GWPPV motif itself (**Fig. 5A**). These residues include aromatic and positively charged amino acids known for strong inter-protein interactions. Additionally, the short-predicted IDR beyond the PB1 domain in IAA7 showed interaction frequencies equal to or greater than the GWPPV motif. Many residues in this disordered region are charged or aromatic, suggesting they may weakly stabilize the IAA7 and ASK1·TIR1 interaction without auxin.

**Fig. 5.**
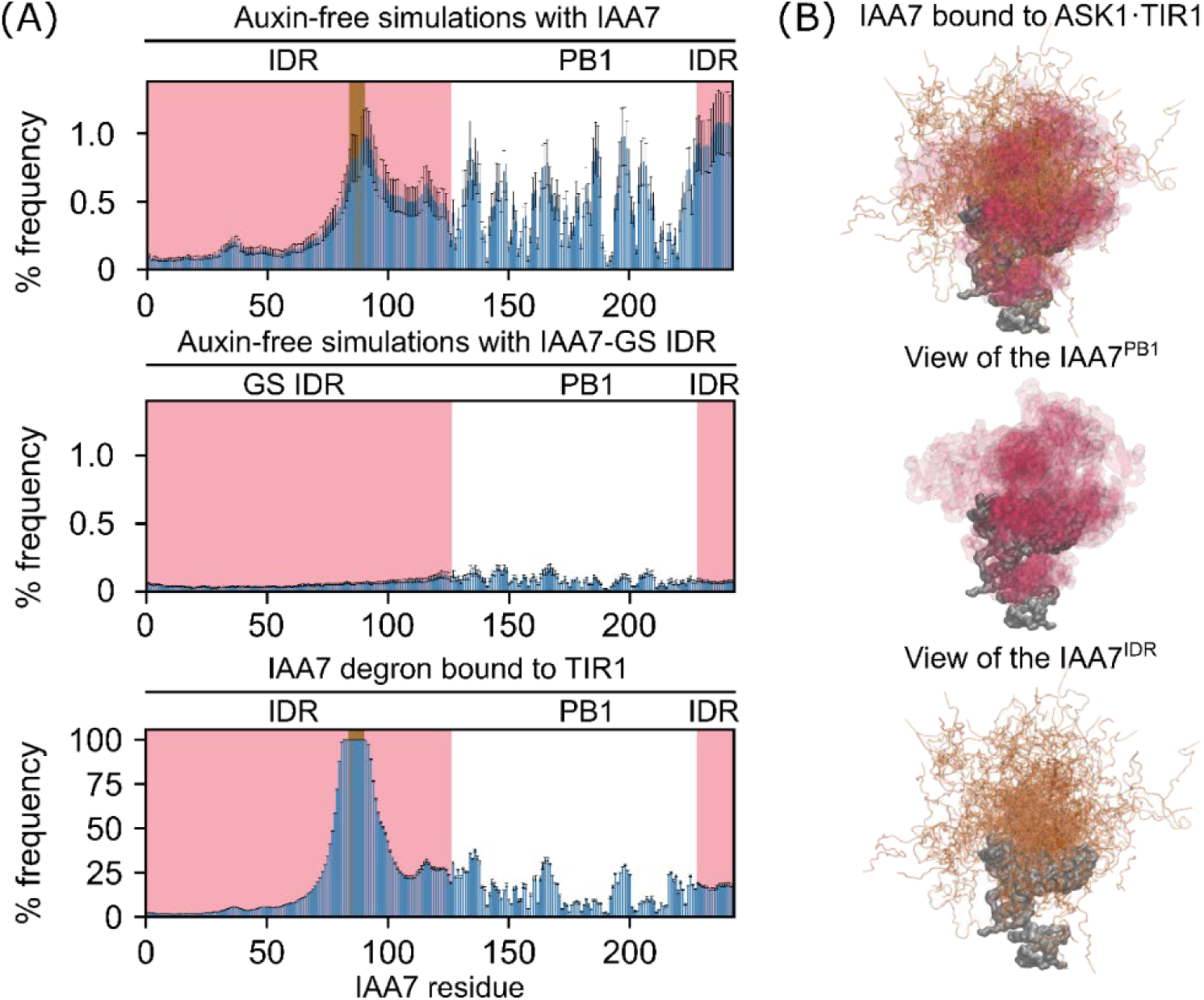
Multiple conformations of IAA7 bound to ASK1·TIR1 shown by CGS. (A) Interaction frequencies of IAA7 with ASK1·TIR1. The percentage of frequency represents the proportion of time that a residue from IAA7 was ≤10 Å apart from ASK1·TIR1 in every simulation frame. In all graphs, the pink shaded area denotes the IDRs, and the brown shaded region represents the GWPPV degron motif. (B) Conformational heterogeneity of IAA7 on the top surface of TIR1 as shown by CGS. Multiple conformational states of IAA7 in the ASK1·TIR1-bound state are achieved due to the flexible regions flanking the degron. Frame overlays are separated, showing only the IDR (orange) or the PB1 (red) for better visualization.

Replacing the entire IDR with poly-GS in IAA7 while keeping the native PB1 resulted in an almost complete loss of the IDR and PB1 interactions with ASK1·TIR1 (**Fig. 5A**). These results indicate that the interaction frequencies between IAA7 and ASK1·TIR1 are specific to the IAA7 sequence and that both PB1 and IDR cooperatively aid in the interaction with ASK1·TIR1.

After identifying regions of IAA7 that drive interactions with ASK1·TIR1 independent of auxin, we next sought to determine inter-residue interactions between ASK1·TIR1 and IAA7 in complex. To simulate the IAA7 in the TIR1-bound state driven by auxin, we fixed the GWPPV core degron motif of IAA7 to its location in the crystal structure of the ASK1·TIR1·auxin·IAA7 complex [21]. The interaction frequency map showed a maximum peak centered on the degron motif, which was expected given that the degron was fixed relative to the auxin binding pocket (**Fig. 5A**). Moreover, the N-terminal region located upstream of the degron exhibited low interaction frequency with ASK1·TIR1, likely due to the IDR structural fluctuations that quickly associate and dissociate across the top surface of TIR1 (**Fig. 5B**). Additionally, the short-predicted IDR located at the end of the PB1 domain showed a substantial binding frequency, supporting an “opened” or expanded IAA7 conformation. This expanded conformation, allows residues distant from the degron motif to interact with the top surface of TIR1 (**Fig. 5A**). When examining different frames of the simulations of the degron tail, the region connecting the degron and the PB1 domain, we observed that the degron tail tethers the PB1, allowing it to bind in different conformations on the top surface of TIR1 (**Fig. 5B**).

Based on our simulations, we propose that during auxin sensing the PB1 domain of IAA7 binds to TIR1 in various conformations while being tethered by the degron tail. Furthermore, the IDR upstream of the degron in IAA7 shows a broad range of expanded conformations (**Fig. 5B**). Overall, in the TIR1-bound state, the top surface of TIR1 acts as a platform, offering numerous sites for transient interactions that may enhance the binding between ASK1·TIR1 and IAA7. This interaction is classified as a fuzzy binding mode, where IAA7 maintains its conformational heterogeneity and establishes multiple contact sites with ASK1·TIR1 [54–56].

## Discussion

Here, we broaden the understanding of the binding modes involved in assembling the auxin receptor complex. Previously, the conformation of IAA7 before binding to TIR1 was unknown, and binding mode models only provided a static view of the conformational ensemble of AUX/IAAs bound to TIR1. Through smFRET, we demonstrated that the average conformation of IAA7 expands in the TIR1-bound state (**Fig. 3B**). This change in the average conformational ensemble of IAA7 is relatively large, with an experimental distance difference of approximately 2.6 nm from the free to the TIR1-bound state at IAA7 C19-C244. Although IAA7 expands, it remains highly dynamic on top of TIR1, as shown by our fluorescence lifetime analysis (**Fig. 4**). This indicates that IAA7 may explore a myriad of expanded conformations during auxin perception, rather than adopting a single expanded conformation as previously thought [36, 37]. Our computational simulations also support this (**Fig. 5B**), showing that the IDR of IAA7 and its PB1 domain establish multiple transient contacts with the top surface of TIR1. We suggest that these additional contacts outside the degron interface potentially support the expanded conformation of IAA7 on top of TIR1 and enhance its affinity for TIR1.

Based on our findings and those of others, the IDR of IAA7 exhibits two distinct binding modes in forming the auxin receptor complex. First, the binding of the degron to the auxin-binding pocket in TIR1 in the presence of auxin can be conceptually classified as a disorder-to-order transition. In this mode, the degron adopts a single, well-defined, and fully ordered binding conformation, with no secondary structure, as shown by X-ray crystallography [21, 54]. However, this disorder-to-order transition is context-dependent, relying on cellular levels of auxin, since IAA7 binds weakly to TIR1 in the absence of auxin [22, 36].

In the second mode, when IAA7 is bound to TIR1, the IDRs upstream and downstream of the degron retain their conformational heterogeneity, sampling both free and TIR1-bound states. These additional transient interactions of the IDR and PB1 with TIR1 are classified as a dynamic “fuzzy” binding mode and contribute to the binding strength between IAA7 and TIR1 [54–56]. From a functional perspective, we interpreted the ASK1·TIR1 and IAA7 binding modes as having two significant outcomes in the context of ubiquitylation. First, the expansion of IAA7 increases the proximity between the lysine acceptor of ubiquitin in IAA7 and the active site of the SCF^TIR1^-E3 ligase [38], thereby enhancing the probability of ubiquitylation (**Fig. 6**). An expanded conformation was also observed in the yeast Cdc4-phosphodegron (CPD) in pSic when bound to the F-BOX protein Cdc4 at the substrate recognition binding site [57]. This suggests that IDR expansion may be a common mechanism for other ubiquitylation targets of SCF-E3 ligases.

**Fig. 6.**
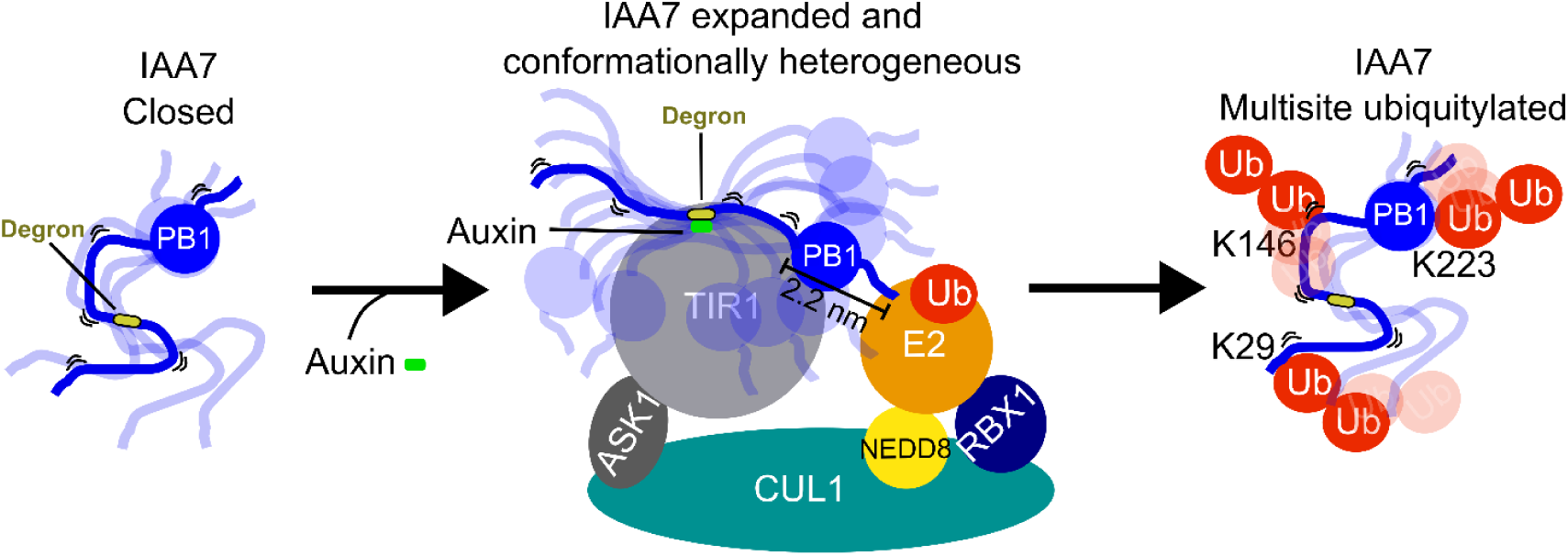
Proposed acrobatics model for IAA7 multisite ubiquitylation by an SCF^TIR1^-E3 ligase. IAA7 is in a “closed” conformation in its free state but tends to adopt expanded conformations within its conformational ensemble. Upon binding to the SCF^TIR1^ complex in the presence of auxin, IAA7 expands exhibiting structural acrobatics. The expansion of IAA7 increases its proximity to the SCF^TIR1^ active site located between the E2∼Ub and TIR1, facilitating ubiquitylation. Because IAA7 remains dynamic and conformationally heterogeneous, different lysine residues enter and exit the SCF^TIR1^ active site, allowing for multisite ubiquitylation in lysine at positions 29, 146, and 223.

Second, the conformational heterogeneity of IAA7 when bound to TIR1 facilitates its multisite ubiquitylation by the SCF^TIR1^-E3 ligase at lysine residues that are spaced far apart. This hypothesis is strongly supported by the fact that IAA7 is ubiquitylated *in vitro* at lysine 29, 146, and 223 [36]. A static interaction of IAA7 with TIR1 would not explain such a spread-out ubiquitylation pattern in a disordered target. Instead, an expanded and conformationally heterogeneous IAA7 would allow different lysine residues in the primary sequence to approach the E3 ligase active site more effectively for ubiquitin transfer (**Fig. 6**). Furthermore, conformational acrobatics should allow the release of the active site as IAA7 is polyubiquitylated, based on *in vitro* assays [36]. In this way, a lysine residue in IAA7 may approach the active site, undergo ubiquitylation, and then be released, allowing another lysine residue to enter the active site for ubiquitylation while IAA7 remains bound to TIR1. This process may resemble an assembly line, where lysine residues in IAA7 are tagged with ubiquitin one after the other while circulating the SCF^TIR1^-E3 ligase active site.

In summary, our results provide novel insights into the functioning of AUX/IAAs IDRs in auxin receptor complex formation and AUX/IAA ubiquitylation. We propose that AUX/IAAs conformational acrobatics is an additional factor determining their binding modes and fate. Thus, the binding versatility and functioning of AUX/IAAs arise from the flexible and dynamic nature of their IDRs.

## Materials and Methods

### Protein purification

ASK1·TIR1 complex was expressed and purified from *Spodoptera frugiperda9* cells as previously described [21] with minor modifications. Briefly, ASK1 was expressed and co-purified with GST-TIR1 using glutathione (GSH) affinity chromatography (SERVA) and anion chromatography (Mono Q™ 5/50 GL). This was followed by TEV tag removal and a final size-exclusion chromatography (SEC) step (Increase 10/300 Superdex 200 pg) using an ÄKTA FPLC system.

IAA7 variants were expressed as GST-tagged proteins in BL21 (DE3) *E. coli* strain. Expression was induced at an OD_600_=1.0 using 1 mM isopropyl β-d-1-thiogalactopyranoside at 16°C overnight. Purification was performed using GSH-agarose beads (SERVA). Briefly, IAA7 cell lysates were incubated with GSH-agarose beads for 30 min, followed by GST tag removal through in-resin thrombin digestion in 20 mM Tris-HCl pH 8.0, 200 mM NaCl. The fractions containing untagged IAA7 variants were pooled and loaded onto an SEC column (HiLoad^TM^ 16/60 Superdex^TM^ 75 pg or 200 pg).

### Protein fluorescent labeling and purification

IAA7 was labeled with ATTO488 and ATTO594 as donor-acceptor pairs following the supplier’s protocol (ATTO-TEC). Untagged, GSH-agarose affinity-purified IAA7 was concentrated to 1-1.5 mM in approximately 250 µL of solution C (20 parts of solution A [phosphate-buffered saline, pH 7.4] and 1 part of solution B [0.2 M sodium bicarbonate, pH 9.0]). The protein solution was mixed with maleimide-ATTO488 (ATTO-TEC) at 0.7 times the molar concentration of the protein and incubated at room temperature for 1 hour. Free ATTO448 dye was removed using cation exchange chromatography with a HiTrap^TM^ SP HP column. Fractions containing IAA7-ATTO488 conjugates were pooled and concentrated to approximately 250 µL in solution C. ATTO594 (ATTO-TEC) was then added at a 1:1 molar ratio relative to the protein, followed by incubation at room temperature for 1 hour. Excess ATTO594 was removed by a second cation chromatography step using a HiTrap^TM^ SP HP column.

### Dynamic and static light scattering (DLS/ SLS)

Simultaneous dynamic and static light scattering was measured at a scattering angle of 90° using a custom-built apparatus equipped with a diode-pumped continuous wave laser (Cobolt Samba 532 nm, Cobolt AB) and a high quantum yield avalanche photodiode, as previously described [58] at 23 °C in 3 mm-pathlength micro-fluorescence cells (105.251-QS, Hellma). Mean scattering intensities from SLS and time-autocorrelation functions of the fluctuations in instantaneous scattering intensities from DLS were recorded over 8 s for at least 100 accumulations. Apparent molecular masses were derived from the mean scattering intensities from SLS measurements, with toluene as a standard as described earlier [58], using a refractive index increment (dn/dc) of 0.188 mL/g.

Translational diffusion coefficients (D) were obtained from the measured autocorrelation functions using the CONTIN algorithm [59], and converted into apparent Stokes radii (*R_S_*) via the Stokes-Einstein equation (*R_S_* = k_B_T/(6πηD), where k_B_ is Boltzmann’s constant, T is the temperature in Kelvin, and η is the solvent viscosity. The refractive index of the buffer was determined with an Abbe refractometer and the viscosity was measured using an Ubbelohde-type viscometer (Viscoboy-2, Lauda). SLS/DLS data were collected for several protein concentrations and the apparent molecular masses and *R_S_* values were extrapolated to zero dilution to obtain the mass and *R_S_* after eliminating the influence of intermolecular interactions.

Compaction indices (CI) were calculated relative to the scaling behavior of intrinsically disordered and globular proteins derived from large sample sets [45, 60] as expressed in scaling laws of type *R_S_= a*M^b^* as described in detail in (45) using the following equation [46, 47]:

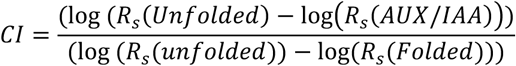

### Confocal fluorescence microscope

Single-molecule fluorescence burst experiments were conducted using a home-built confocal microscope equipped with a pulsed fiber laser (FemtoFiber pro TVIS, Toptica Photonics) operating at 488 nm and a repetition rate of 80 MHz. The pulses were synchronized with a diode-based laser (LDH P-C-485B, Picoquant) operating at 485 nm and 20 MHz. A single-mode fiber (LMA-8, NKT Photonics) was used for spatial filtering, and a 60X microscope objective (UPlanApo 60x/1.20W, Olympus) was used for excitation and fluorescence light collection. Dichroic beam splitters (ZT405/488/594/647rpc, ZT594rdc, Chroma Technology Corp.) and a polarizing beam splitter (CVI Laser Optics) were used to split the emission light and guide it onto single-photon avalanche diodes (SPCM-AQRH-14-TR, Excelitas Technologies Corp.) which serve as confocal pinholes. Spectral filters were used to set the spectral range for the donor channel (LP496, BP25/50) and the acceptor channel (LP615, BP629/56), all from Semrock Inc (IDEX Corp.). Pulses from the detectors were fed into a TCSPC board (MultiHarp 150, Picoquant) operating in time-tagged time-resolved mode with 80 ps time resolution.

### Single-molecule FRET measurements and fluorescent lifetime analysis

The measurements were carried out with 50-100 pM of doubly-labeled IAA7 alone or preincubated with 10 µM indole-3-acetic acid (IAA, auxin) and/or 1 µM of ASK1·TIR1 for 20 minutes. For burst experiments, a pulsed-interleaved excitation scheme was used, with the donor molecule excited at 80 MHz and 50 µW (488 nm) and the acceptor at 20 MHz and 10 µW (594 nm). Detected photons were sorted by their arrival times and with respect to a synchronization signal of 20 MHz. Fluorescence bursts were identified by applying a threshold criterium to a sliding photon density average with consecutive positively selected photons combined into bursts. For single-burst FRET analysis, two threshold criteria were used - one threshold accounted for the integrated emission after donor excitation and one threshold for the emission upon direct excitation of the acceptor - to select for molecules bearing both a donor and acceptor. The individual photon counts were corrected for background, spectral cross talk (α), different quantum yields and detection efficiencies of donor and acceptor molecules (β), differences in excitation intensities and absorption cross-section (γ) and direct excitation of the acceptor at the donor excitation wavelength (δ). This led to an individual energy transfer (*E*) [61]:

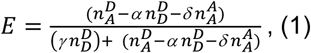

where 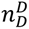 and 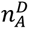 are the background-corrected number of detected donor and acceptor photons after donor excitation, and 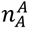 is the background-corrected number of detected acceptor photons after acceptor excitation. In addition, the labeling stoichiometry ratio (S), was determined accordingly:

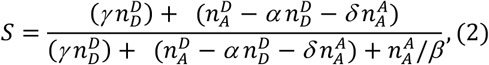

This procedure allowed us to filter out bursts originating from molecules lacking an active acceptor. Finally, the mean FRET efficiencies were determined from the *E*-histograms by fitting one or multiple Gaussian functions to determine the mean energy transfer value of each distribution. Distance averaging due to conformational flexibility during transit through the focus was considered using the integral:

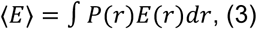

where *E*(*r*) = 1/(1 + (*r*/*R*_0_)^6^) is the distance dependence of the energy transfer, and *P*(*r*) is an excluded-volume probability distribution of the dye-to-dye distance given by

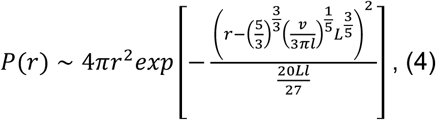

The values *v* and *l* were deduced from the distance distribution by EOM modeling of the X-ray scattering profile of IAA7, with *v*=0.01 and *l* =6.7A (see X-ray scattering section). The parameter L describes the contour length, with L=Nb, where N is the number of residues between the fluorescent markers and b is the distance between two Cα atoms (b=0.38 nm). Finally, the obtained values of *R_ee_^FRET^* were corrected for the lengths added by the linker to which the dye molecules are attached, considering their combined length of 10.7 b, in total: *N* = *N_protein_* + *N_linker_*.

Fluorescence lifetimes of the donor molecule in a doubly labeled sample (*τ_DA_*) were estimated from the mean relative arrival times of all donor burst photons. The fluorescence lifetimes were normalized to the fluorescence lifetime in the absence of any acceptor molecule (*τ_D_*), and plotted against corresponding transfer efficiencies in two-dimensional scatter plots. For a fixed distance between the donor and acceptor, the ratio *τ_DA_*/*τ_D_* must equal (1-*E*), leading to a diagonal line. For systems that rapidly sample a distance distribution (equation 3), this ratio significantly deviates from this line [52].

### Small-angle X-ray scattering (SAXS)

X-ray scattering experiments were performed in transmission mode using a SAXSLAB laboratory setup (Retro-F) equipped with an AXO microfocus X-ray source. The AXO multilayer X-ray optic (AXO Dresden GmbH) was used as a monochromator for Cu-*K*_*α*_radiation (*λ* = 0.154 nm). A two-dimensional detector (PILATUS3 R 300K; DECTRIS) was used to record the 2D scattering patterns. SAXS experiments were conducted using refillable capillaries with an outer diameter of 1 mm (BioSAS JSP stage, SAXSLAB/Xenocs). The intensities were angular-averaged and plotted versus the scattering angle (*q*). The measurements were performed at room temperature and corrected for background, transmission, and sample geometry. The measurement times were 10 hours. Subsequent EOM analysis (GAJOE - version 2.1) using default parameters - a pool size of 10,000 conformations generated in disordered mode, an ensemble size of a maximum of 20 conformations, and 100 iteration cycles revealed the most probable distance distributions [62, 63].

### Simulation of IAA7 SAXS profiles and reweighting

SAXS profiles were simulated for molecular dynamic (MD) trajectories and reweighted to improve agreement with experimental SAXS data. SAXS profiles were generated for individual frames using the autoSCTR tool. To reduce computational complexity, the simulation was downsampled to 1,000 evenly spaced frames, capturing the structural diversity of the trajectory while maintaining computational efficiency. The SAXS profiles from all sampled frames were merged to construct a composite profile representing the simulation. To ensure comparability with experimental SAXS data, the q-values of the simulated SAXS profiles were interpolated to match the range observed in experimental datasets, accounting for variations introduced by beam source, detector, and other experimental parameters. The BME package (https://github.com/KULL-Centre/BME) was used to reweight the simulated SAXS profiles to align more closely with experimental data. The BME.Reweight function applied a Bayesian Maximum Entropy framework to adjust the contributions of individual frames, optimizing agreement between simulated and experimental SAXS profiles. Details on this methodology are available in the BME GitHub repository and the associated protocol [51].

### Coarse-grained simulation of IAA7 bound to ASK1·TIR1

Simulations were carried out using the LAMMPS simulation engine with default Mpipi parameters [50]. The Mpipi model is a coarse-grained, one-bead-per-residue model optimized for simulating intrinsically disordered regions (IDRs). The primary reason for using one-bead-per-residue simulations was the large size of the system simulated (754 amino acids for ASK1·TIR1 and 243 residues for IAA7). In our simulations, the IDR was modeled as flexible chains, while folded domains were modeled as fixed rigid bodies. To convert atomistic structures to coarse-grained models, the location of the α-carbon atoms was used to place each bead for each amino acid. The IAA7 PB1 structure was predicted using AlphaFold2 and ColabFold with the following specifications: structures were ranked by pLDDT score, max_msa was set to 512:1024, use_turbo was set to True, num_models was set to 5, use_ptm was set to True, num_ensembles was set to 8, max_recycles was set to 3, is training was set to False, and num_samples was set to 1. For the predicted IAA7 IDR, simulations started with non-overlapping amino acids. The ASK1·TIR1 structure was based on a previously reported crystal structure [21]. Missing amino acids were filled in by generating a structure of ASK1·TIR1 using ColabFold with the same parameters as for IAA7. The AlphaFold2-generated structure was aligned with the crystal structure in VMD, and the coordinates of the missing amino acids were saved and included in the final simulations. All simulations were conducted in a 500Å-by-500Å box with periodic boundary conditions at a temperature of 300 °K. For the “auxin-free” simulations, twenty replicates were carried out to reduce variability from the random placement of IAA7 and ASK1·TIR1 in the box. Each replicate consisted of 100,000,000 steps at 10 femtoseconds per step. The first 500,000 steps of each simulation were discarded to allow for equilibration, resulting in 99,500,000 steps used for analysis. Coordinates from the simulation were saved at intervals of 10,000 steps. For the ASK·TIR1-bound simulations, the IAA7 GWPPV motif was fixed in the same position as in the ASK1·TIR1·auxin·IAA7 crystal structure [21]. Apart from these 5 residues, all other IAA7 IDR residues were allowed to move freely. For the auxin-bound simulations, due to minimal variability in the relative position of IAA7 to ASK1·TIR1, three replicates were carried out. Each replicate consisted of 200,000,000 steps at 10 femtoseconds per step. Like the auxin-free simulations, the first 500,000 steps were discarded for equilibration, and coordinates were recorded every 10,000 steps. For all simulations, the Python package SOURSOP was used for analysis, which is based in part on MDTraj [64, 65]. For contact frequency calculations, residues were considered “in contact” if they were less than 10 Å apart. Determination of residue contact was binary; if a residue was within at least one residue of the other protein, it was considered “in contact” for the calculation, as determined using the compute_contacts function in MDTraj.

## Acknowledgments

We thank Micheal Niemeyer for his support and advice, and Susanne Freund for technical assistance in the DLS/SLS experiments. This work was supported by the Deutsche Forschungsgemeinschaft (Research Training Group RTG2467), and core funding of the Leibniz Institute of Plant Biochemistry (IPB).

## Supporting Text 1. Liquid chromatography-mass spectrometry (LC-MS) for labeling detection

Fluorescently labeled IAA7 was separated by SDS-PAGE, and the corresponding band was excised and subjected to in-gel digestion with either Trypsin, or GluC, or both. Extracted and dried peptides were desalted, dissolved in 5% acetonitrile, 0.1% trifluoroacetic acid, and injected into an EASY-nLC 1200 LC-system (Thermo Fisher Scientific). Peptides were separated using C18 reverse-phase chromatography, and the eluted peptides were electrosprayed online into a Fusion Lumos Tribrid mass spectrometer (Thermo Fisher Scientific). The spray voltage was 2.0 kV and the capillary temperature 305°C.

A full MS survey scan was carried out with a chromatographic peak width set to 15 s, a resolution 60,000, automatic gain control (AGC) set to standard, and a maximum injection time (IT) of 100 ms. MS/MS peptide sequencing was performed using a parallel reaction monitoring (PRM) scan strategy (without retention time scheduling) with HCD fragmentation containing *in silico* generated target peptide m/z ratios according to the lists in Supporting Tables SI and S2. The top 15 MS/MS scans were acquired in the Orbitrap with a resolution 15,000, mass-to-charge ratios (m/z) between 150 and 2000, AGC target set to 300%, Maximum IT 22 ms, isolation width 1.6 m/z, and normalized collision energy 28%.

MS/MS spectra were used to search the TAIR10 database (ftp://ftp.arabidopsis.org) amended with the protein sequence of the modified IAA7 protein by the Mascot software v.2.7 linked to Proteome Discoverer v.2.1. The enzyme specificity was set to the respective enzyme(s) and up to five missed cleavages were tolerated. Carbamidomethylation of cysteine was set as a fixed modification, while oxidation of methionine and ATTO488 or ATTO594 modification of cysteine were set as variable modifications. The precursor tolerance was set to 10 ppm and the product ion mass tolerance was set to 0.02 Da. A decoy database search was performed to determine the peptide spectral match (PSM) and peptide identification false discovery rates (FDR). The ptmRS module was utilized to map ATTO488 or ATTO594 labeling sites within the primary sequence of IAA7, employing standard settings.

**Fig. S1.**
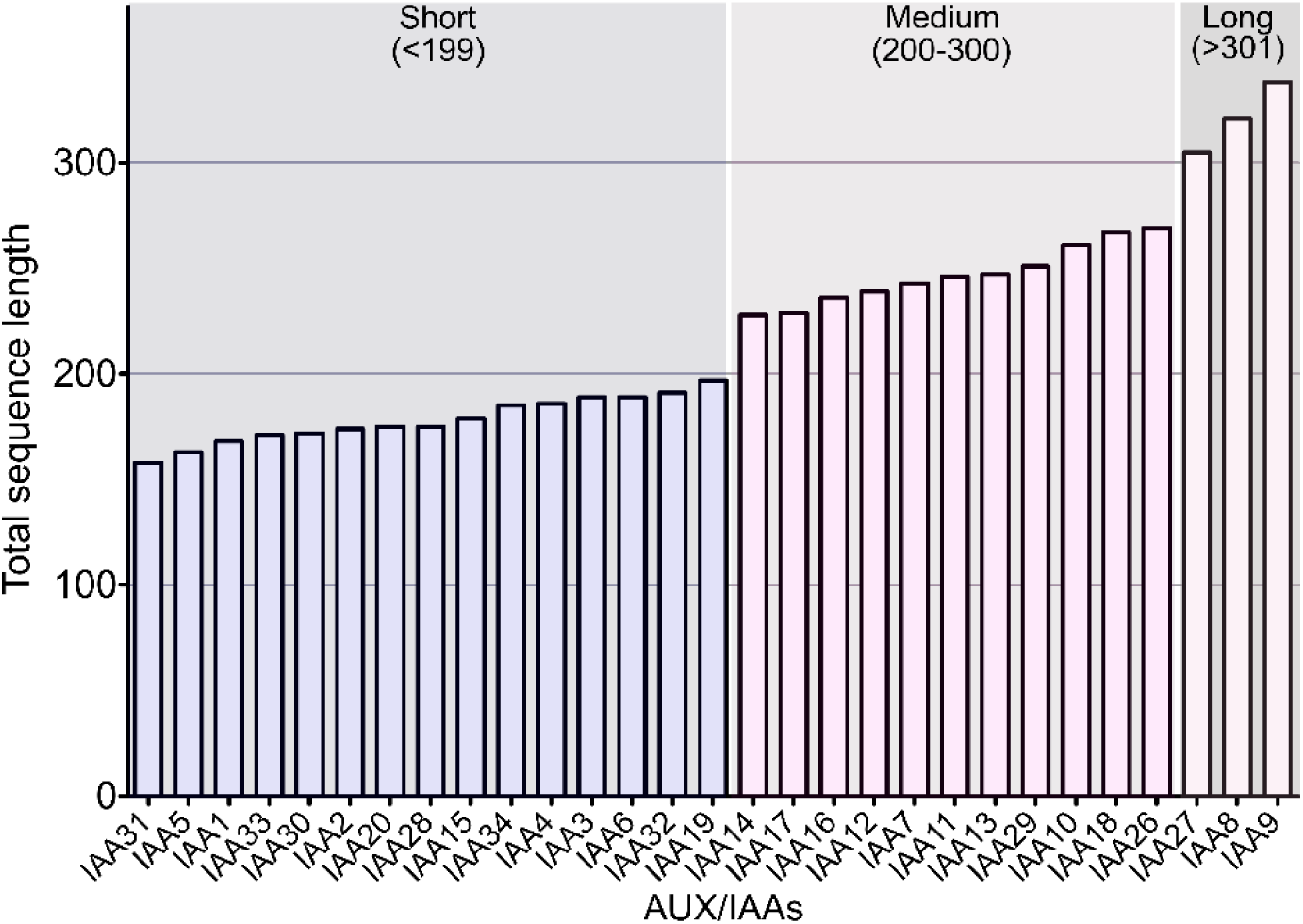
*Arabidopsis thaliana* AUX/IAAs grouped by sequence length. The 29 AUX/IAAs were arbitrarily grouped based on the total number of residues into short (<199 residues), medium (200-300 residues), and long (>300 residues).

**Fig. S2.**
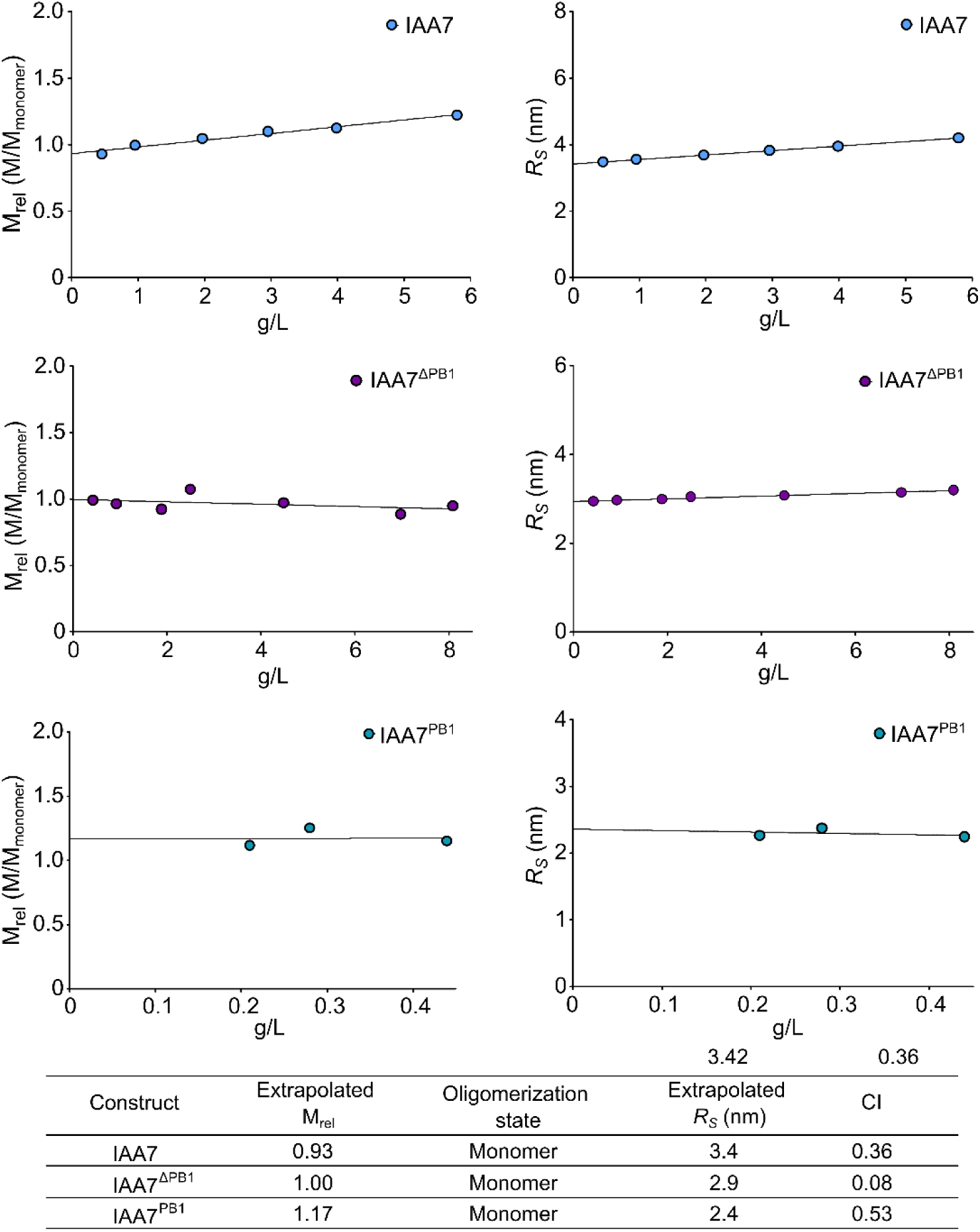
Apparent relative masses and *R_S_* determined by DLS/SLS. Measurements were performed in serial dilutions, and results were extrapolated to zero protein concentration to determine the relative masses (measured mass divided by the theoretical mass of the monomer), reflecting the average oligomeric state, and *R_S_*.

**Fig. S3.**
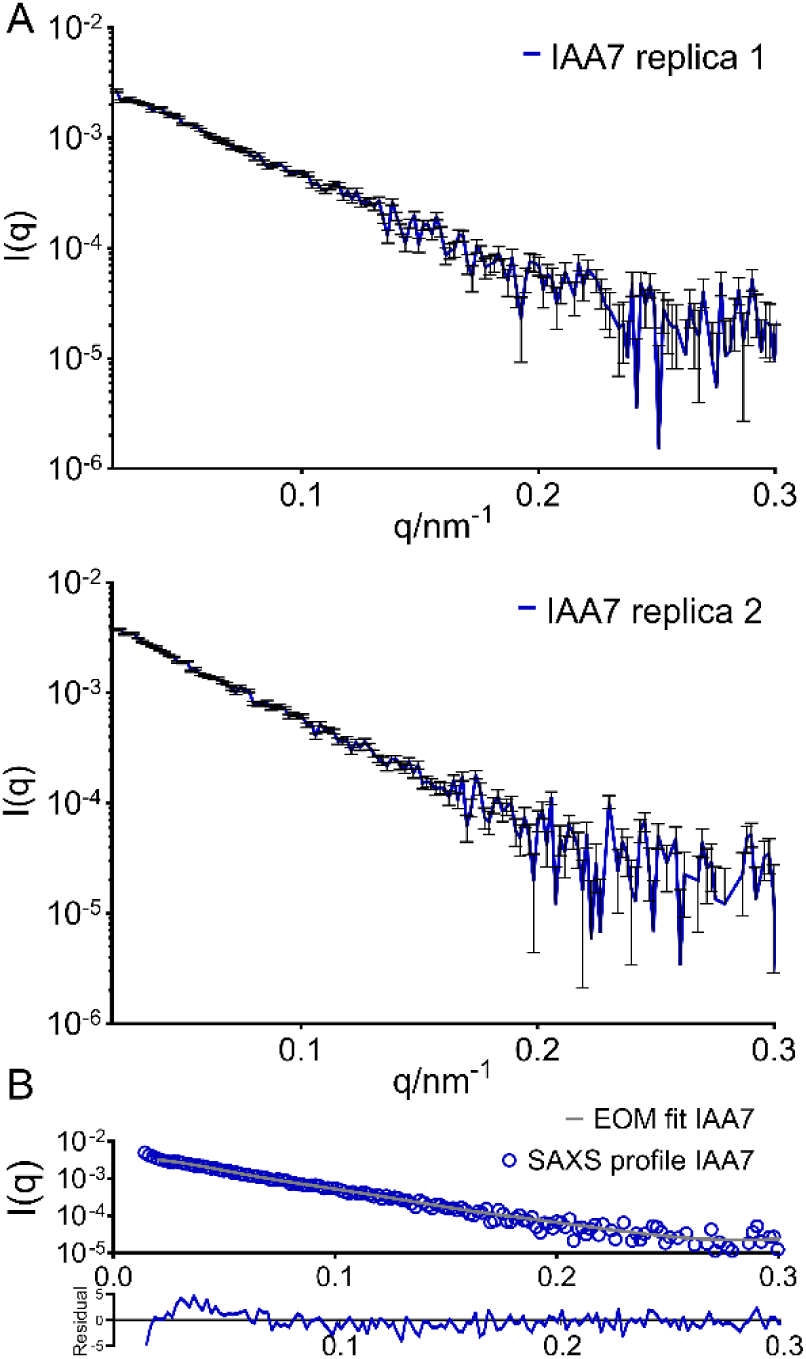
SAXS profile and ensemble optimization method (EOM) analysis of IAA7. (A) SAXS profiles were recorded in two technical replicates with a protein concentration of 2 g/L for IAA7. In each replicate, the standard deviation of I(q) is shown in black bars. (B) The SAXS profiles of IAA7 were analyzed using EOM, with the analysis performed on the mean scattering intensities obtained from two independent replicas. The initial pool contained 10,000 models with random conformations, from which those that best matched the experimental SAXS average profile were selected by the EOM procedure.

**Fig. S4.**
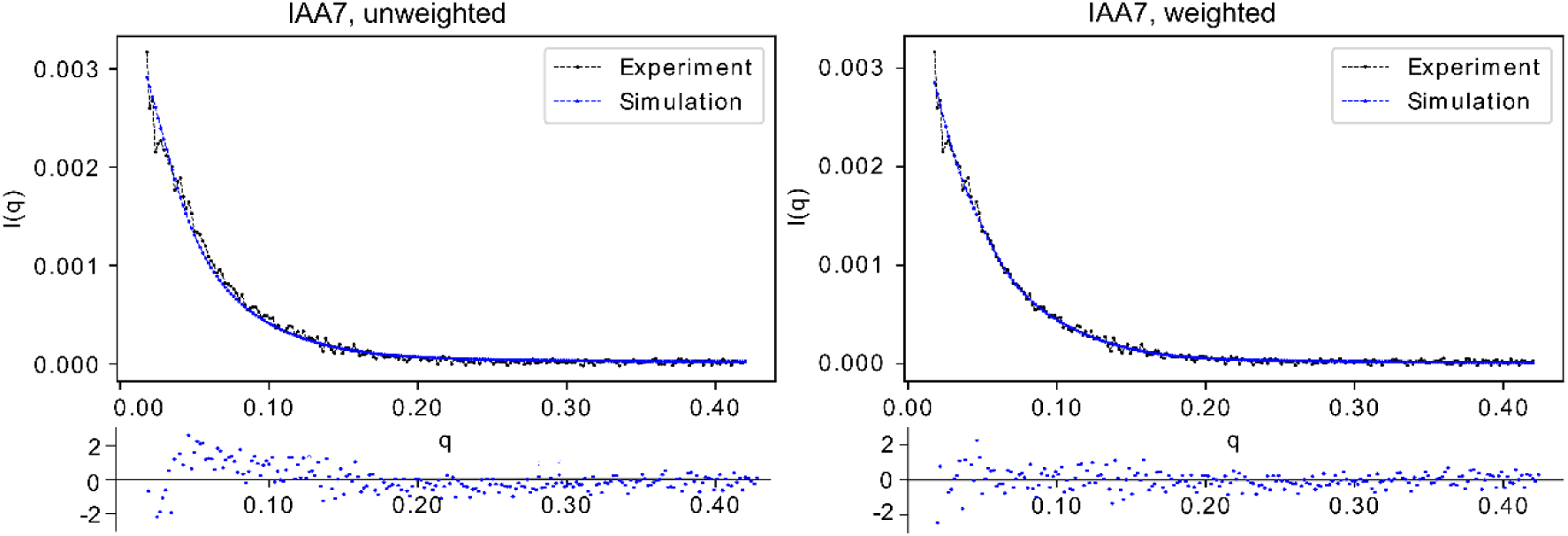
Simulated SAXS curve of IAA7. SAXS profiles were simulated for MD trajectories and reweighted to enhance agreement with experimental SAXS data. Individual frames were processed using the autoSCTR tool to generate SAXS profiles. The BME package (https://github.com/KULL-Centre/BME) was used to reweight the simulated SAXS profiles, ensuring closer alignment with experimental data.

**Fig. S5.**
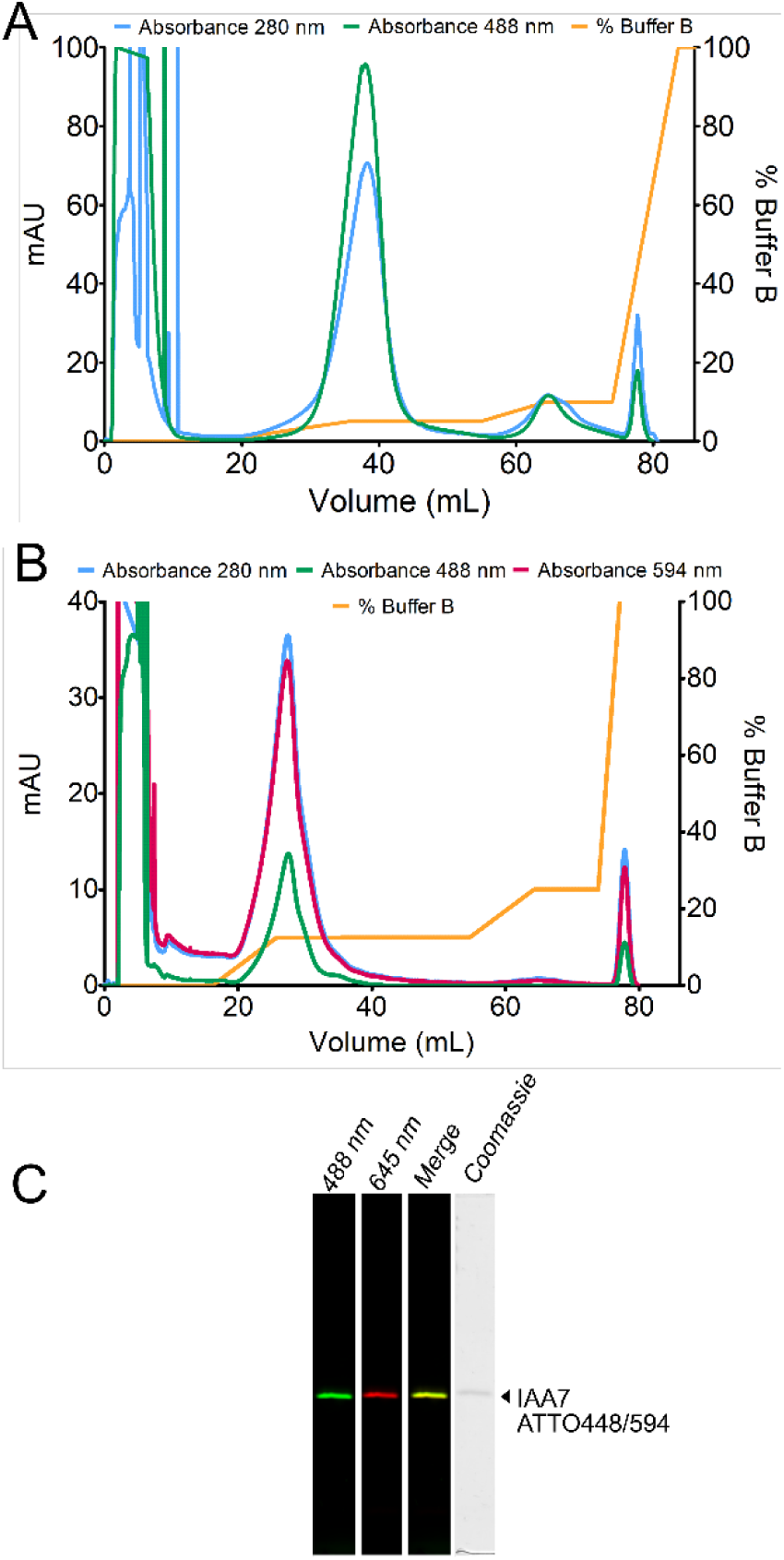
Elution profile of single and double-labeled IAA7. (A) IAA7 was initially labeled with ATTO488 and purified using a HiTrap^TM^ SP HP column with an ÄKTA FPLC system. (B) IAA7-ATTO488 conjugates were next labeled with ATTO594 and purified using the same column and system. In both cases, IAA7 was eluted in 20 mM potassium phosphate buffer (2.76 mM K_2_HPO_4_ and 17.24 mM KH_2_PO_4_), pH 6.0 with a gradient from 0.02 to 1 M of NaCl. Wavelength detection at 280 nm, 488 nm, and 594 nm were used to identify the elution of protein, ATTO488 and ATTO594, respectively. (C) Fluorescent gel scan detection of doubled-labeled IAA7 fractions in Typhoon FLA 9500 shows fluorescence emission of the respective dyes.

**Fig. S6.**
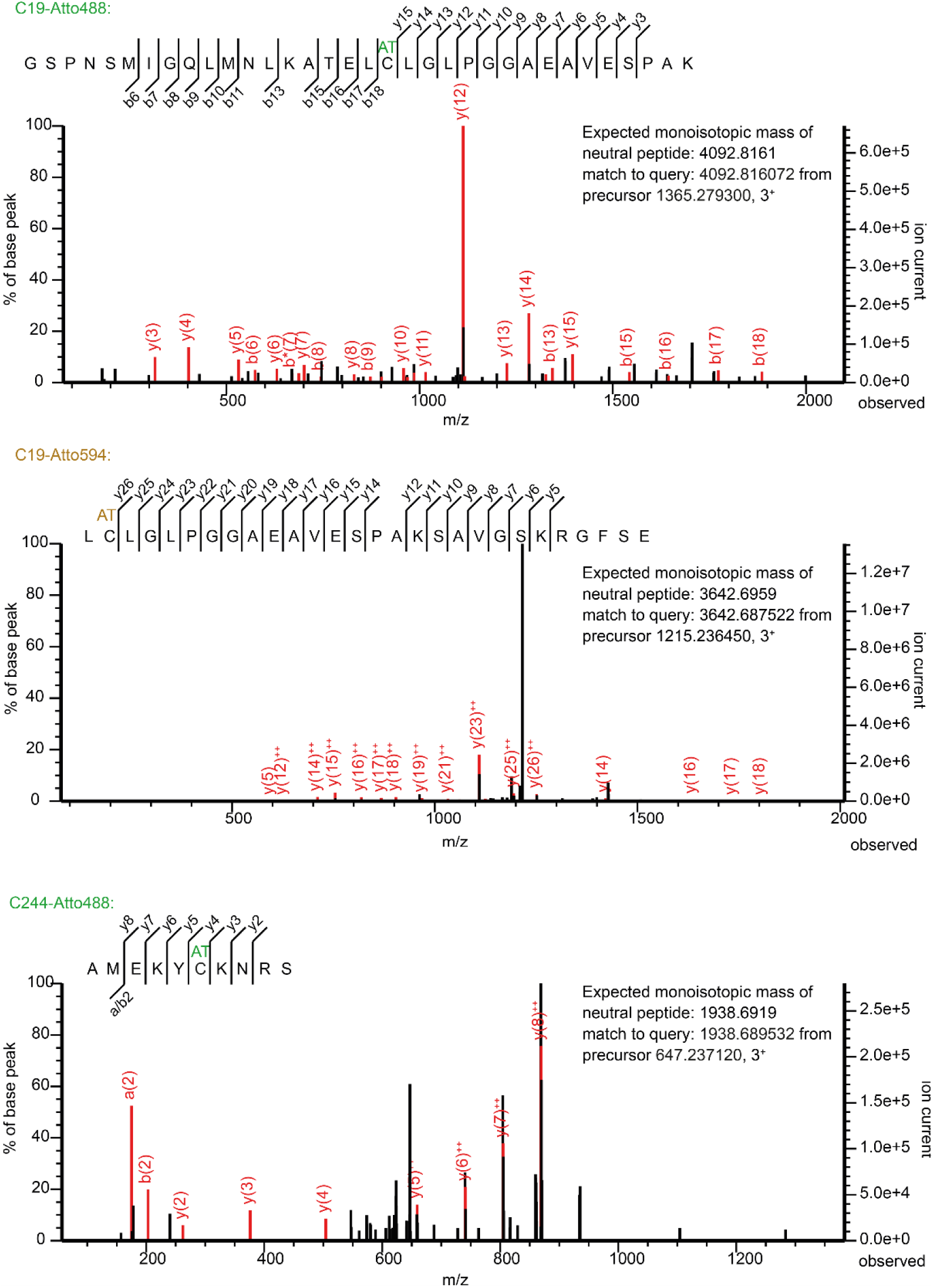
LC-MS/MS identification of IAA7 labeled with ATTO488 and ATTO594. MS/MS fragmentation patterns corresponding to ATTO-labeled IAA7 peptides at C19 or C244 were identified through targeted LC-MS/MS-analysis of purified proteins. ATTO488 modification was detected at both C19 and C244, while ATTO594 was identified only at C19 due to low peptide coverage at the C-terminus. AT: ATTO modification.

**Fig. S7.**
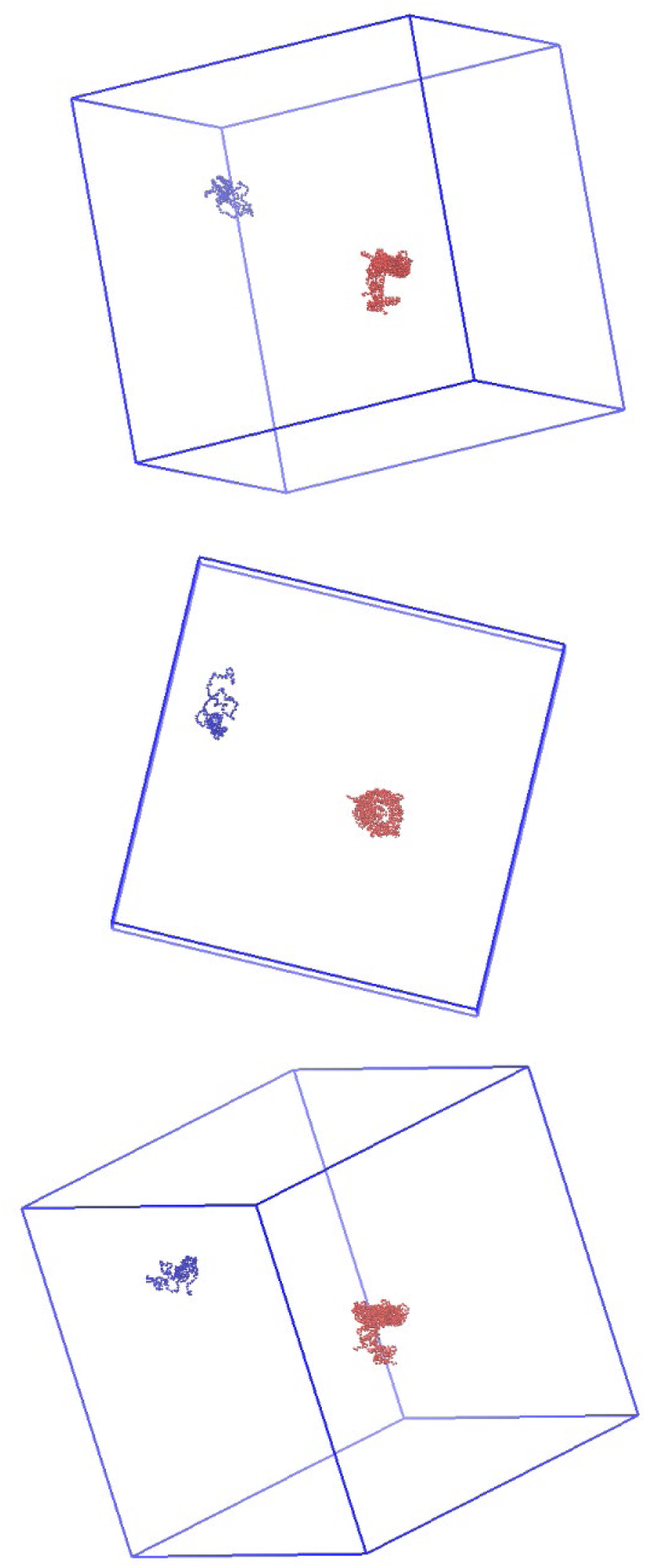
Box space of 500Å-by-500Å for determination of IAA7 and TIR1 interaction frequency. Coarse-grain simulations (CGS) were used to determine the interaction frequency between IAA7 and TIR1 in the auxin-free state. IAA7 is depicted in blue, while TIR1 is shown in orange. The rendering was performed using Visual Molecular Dynamics (VMD).

**Table S1.**
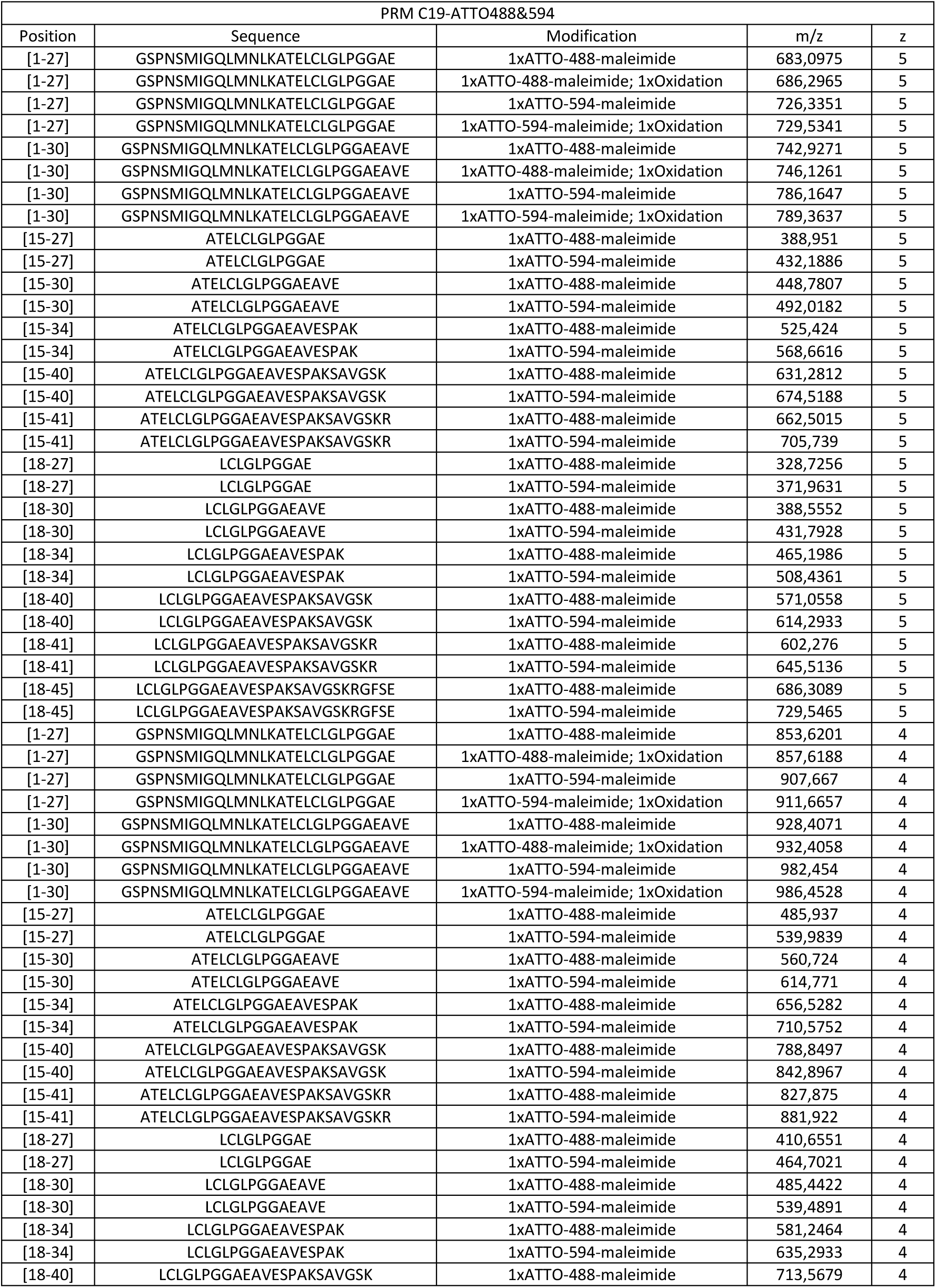
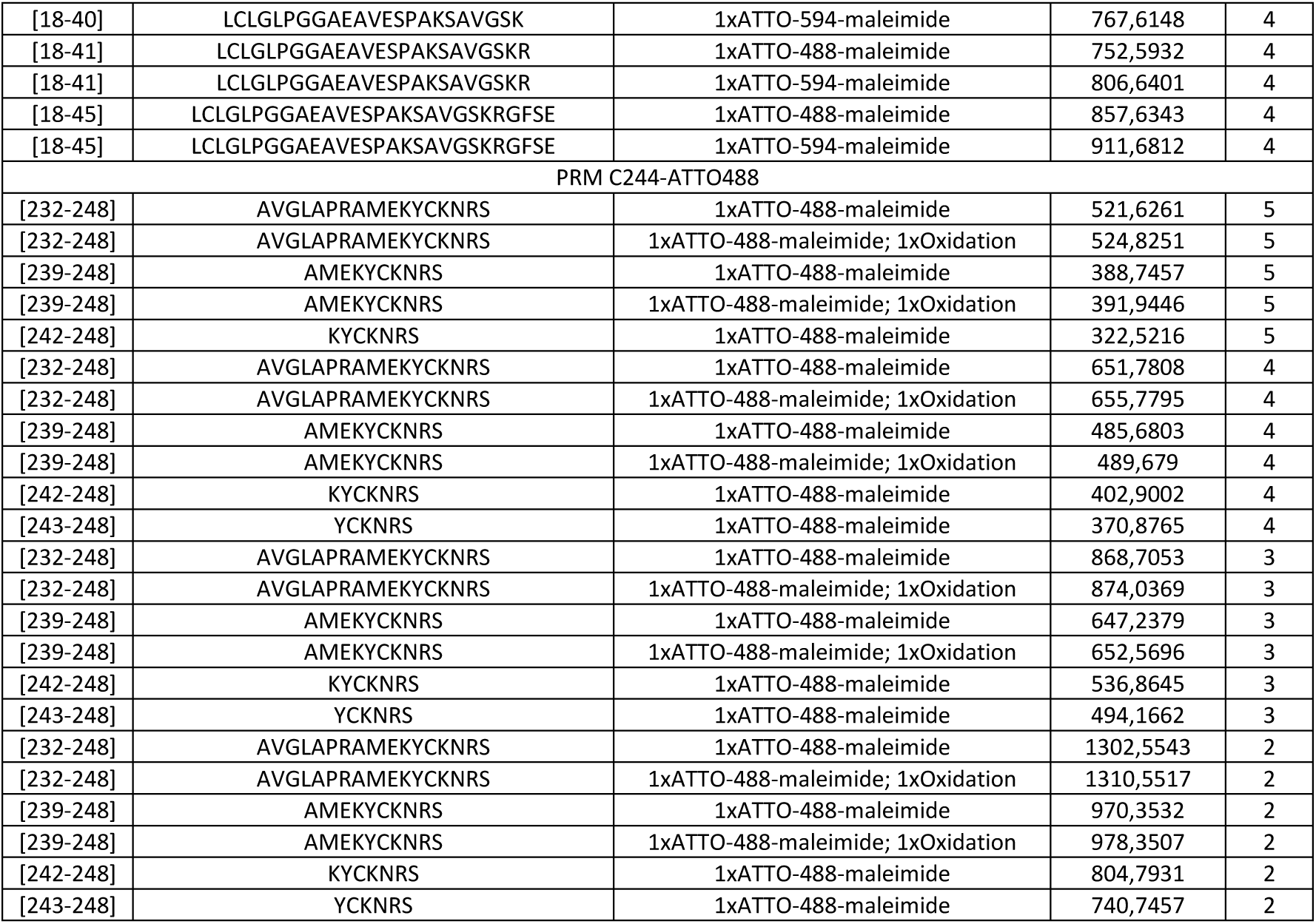
List of target peptides for parallel reaction monitoring (PRM) analysis with ATTO dyes labeling.

**Table S2.** smFRET fitting parameters.

| | Replicas | $\langle E \rangle$ | Fit Error of the mean | Gaussian width | Gaussian Sigma | $R_{ee}$ (nm) linker corrected |
| --- | --- | --- | --- | --- | --- | --- |
| IAA7 | 3 | 0.38 | 0.002 | 0.19 | 0.13 | 7.2 |
| IAA7+auxin | 3 | 0.40 | 0.002 | 0.19 | 0.13 | 7.0 |
| IAA7+ASK1·TIR1 | 3 | 0.39 | 0.003 | 0.27 | 0.19 | 7.1 |
| ASK1·TIR1·IAA·IAA7 | 3 | 0.21 | 0.003 | 0.19 | 0.12 | 9.6 |
|  |  | 0.39 | Fixed | 0.19 | 0.13 | 7.1 |

